# COSMOS: COmmunity and Single Microbe Optimisation System

**DOI:** 10.1101/2024.10.31.621216

**Authors:** Lavanya Raajaraam, Karthik Raman

## Abstract

Bioprocessing harnesses the potential of microbial communities and monocultures to convert renewable resources like agricultural by-products and wastewater into valuable products. While monocultures offer simplicity and control, microbial communities provide metabolic diversity, enabling complex substrate breakdown and cooperative biosynthesis. However, balancing efficiency with stability requires a nuanced approach, as each system presents unique advantages and limitations, particularly under different environmental conditions. To systematically explore the potential of these systems, we developed the COmmunity and Single Microbe Optimisation System (COSMOS), a dynamic computational framework that compares microbial monocultures and communities across diverse fermentation conditions. COSMOS identifies optimal microbial systems tailored to specific carbon sources and environments. Our findings highlight the impact of key factors such as environment, microbial interactions, and carbon sources on the biosynthetic capabilities of both communities and monocultures. A key result is the identification of the *Shewanella oneidensis* – *Klebsiella pneumoniae* community as the most efficient producer of 1,3-propanediol under anaerobic conditions. Notably, this aligns with previous experimental findings, with COSMOS accurately predicting the optimal carbon source concentration and inoculum ratio used. Additional findings highlight the value of communities for nutrient-limited processes and emphasise the importance of computational screening to balance productivity with ease of control. The insights gained from this study offer a roadmap toward optimising microbial systems for sustainable bioprocesses and circular bio-economies.

**Importance:** The transition to sustainable biomanufacturing is vital for reducing reliance on fossil-based resources in industries such as biofuels, wastewater treatment, and pharmaceuticals. While monocultures are preferred for their predictability, microbial communities offer greater metabolic versatility, particularly in nutrient-limited environments. However, their complexity and the lack of systematic evaluation tools have hindered wider adoption. Identifying the most efficient microbial system—whether monoculture or community—and optimizing it for resource efficiency is key to sustainability and aligns with the UN Sustainable Development Goals (SDG 9 and SDG 12). COSMOS addresses this gap by providing a computational framework to assess microbial systems, minimizing the need for extensive experimental testing. By enabling data-driven selection of scalable and efficient bioprocessing strategies, COSMOS supports the transition to circular bio-economies and sustainable manufacturing, ensuring resource-efficient production while reducing environmental impact.

## 1 Introduction

Biomanufacturing supports sustainable development by converting renewable resources, such as agricultural waste or wastewater, into valuable products like biofuels, pharmaceuticals, and bioplastics (Süntar et al. 2021). Both microbial monocultures and communities are used in these processes, each with distinct strengths and challenges. Monocultures are easier to control, manipulate, and engineer for specific product yields, making them ideal for simple, well-characterised bioprocesses. However, their productivity can reach a plateau, even after genetic optimisation (Pachapur et al. 2015).

Microbial communities, on the other hand, leverage complementary metabolic capabilities and interspecies cooperation (Ibrahim, Raajaraam, and Raman 2021). This has sparked significant interest in engineering co-cultures tailored to achieve specific bioprocess objectives (Rafieenia et al. 2024). Notably, even communities composed of strains from the same organism can outperform individual strains. (H. Peng et al. 2024) demonstrated this by developing a molecular toolkit of auxotrophic and overexpression yeast strains, enabling the construction of diverse two- and three-member communities with distinct metabolic capabilities. In contrast, heterologous communities, while more complex, offer unique advantages. In these systems, organisms break down complex substrates—such as lignin or cellulose—into simpler compounds, facilitating resource-sharing (X. “Nick” Peng, Gilmore, and O’Malley 2016; Raajaraam and Raman 2024). Communities are also less prone to feedback inhibition because one species may utilise by-products of another, leading to enhanced growth and stability (Ghosh, Chowdhury, and Bhattacharya 2016). Furthermore, the ‘division of labour’ across species helps distribute the metabolic burden, improving production efficiency for complex products (Rafieenia, Atkinson, and Ledesma-Amaro 2022). However, communities are inherently harder to manage and require careful optimisation to maintain stability and maximise productivity.

Given these trade-offs, the choice between monocultures and communities is dependent on the nature of the bioprocess. The cooperative behaviour of communities provides resilience and efficiency (Ziganshin et al. 2013), whereas monocultures are well-studied and can be easier to manipulate. Designing effective bioprocesses involves identifying the most suitable microbial system for specific substrates or desired products. The decision is not always as straightforward as selecting communities for lignocellulosic biomass conversion and monocultures for simpler fermentations (Liang et al. 2020). Selecting the optimal microbial system for other processes can be challenging, as it would require testing numerous combinations of organisms and environmental conditions—an approach that is both labour-intensive and impractical (Fischer, Klein-Marcuschamer, and Stephanopoulos 2008). Therefore, computational algorithms become essential for managing this complexity and streamlining the selection process (García-Jiménez, Torres-Bacete, and Nogales 2021).

While some algorithms optimize specific co-cultures, few can systematically evaluate multiple microbial systems to identify the most suitable option for a given environment. For instance, (Martinez et al. 2022) have developed an algorithm to optimise substrate-pulsing to obtain stable communities. Other studies have explored strategies to induce microbial communities for specialized product synthesis, with various computational approaches developed to regulate these processes (Boruta 2023). There are also studies like (C. Foster et al. 2021) which focus on a single coculture and model growth kinetics to study how interspecies cell fusion can influence bioproduction. Other algorithms like FLYCOP (García-Jiménez, García, and Nogales 2018) optimises a specific microbial co-culture for synthesizing the desired product. While (Wilken et al. 2018), use constraint-based modelling to find suitable bacterial partners for anaerobic fungi in co-cultures. However, these tools are limited in scope—they cannot comprehensively analyse multiple communities or compare them with monocultures to determine the most effective system.

In this work, we present COSMOS, an algorithm that overcomes these limitations and identifies the best microbial system by building pairwise communities from a set of individual organisms. COSMOS integrates dynamic Flux Balance Analysis (FBA) (Henson and Hanly 2014; Mahadevan, Edwards, and Francis J Doyle 2002) and Flux Variability Analysis (FVA) to simulate the growth of communities and their constituent monocultures (Gudmundsson and Thiele 2010). Moreover, it compares the performance of each community against the highest-performing monoculture, rather than relying on the average productivity of the monocultures, ensuring that the best microbial system is chosen. It also enables users to make informed decisions not only based on productivity but also on additional factors, such as the abundance distribution of the community or any biological insights that might favour one community over another.

In summary, this work contributes to the development of sustainable biomanufacturing by identifying when communities or monocultures should be used and how bioresources can be efficiently employed. It marks a step forward in creating economically viable, eco-friendly processes, aligning with the global effort to promote sustainable industrial practices. This framework aligns with the growing need for optimised bio-based processes that reduce waste and reliance on fossil resources, promoting a circular economy.

## 2 Methods

### 2.1 Flux Balance Analysis (FBA)

The metabolic network of the organism is represented as a stoichiometric matrix *A* of size *m* × *n* where *m* is the number of metabolites and *n* is the number of reactions. The entries in each column represent the stoichiometric coefficients of the metabolites that constitute the reaction. The linear programming (LP) problem is denoted by

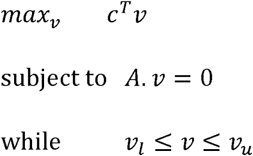

Where *c* is a vector of weights denoting the contribution of each reaction to the objective function, *v*@@@ *R^n^* is the vector of metabolic fluxes, *v_l_*, and *v_u_* are the lower and upper bounds, respectively.

### 2.2 Dynamic FBA

Dynamic FBA is performed using the static optimisation approach where the entire batch time is divided into intervals, and the LP is solved at each time interval as a standard FBA problem. While FBA assumes that the cell maintains an intracellular steady state at a given time point, all (μ) and intracellular fluxes (*v*), including product secretion rates (*v_p_*) at any given time point variables—both intracellular and extracellular—are subject to time variations. The growth rate are determined by solving a standard FBA problem at the biomass concentration *X*. Instead of using fixed substrate uptake rates as in the case of classic FBA, we utilise extracellular (*v_s_*) based on specific kinetic parameters, as shown in Figure 1. These rates reflect the concentrations of substrates (S) and products (P) to calculate dynamic substrate uptake rates maximum uptake capabilities at each time point and are applied as constraints in the calculations.

**Fig 1.**
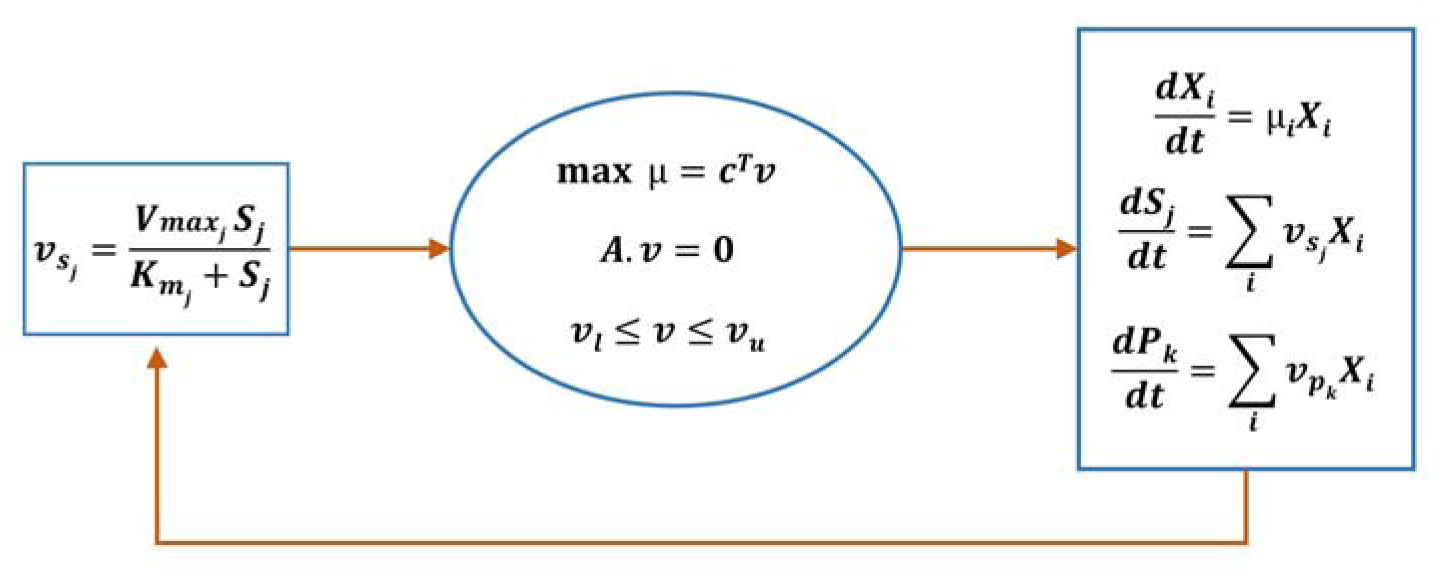
Dynamic flux balance analysis model for a microbial community. The substrate flux 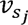 for metabolite *j* is determined using substrate concentration *S_j_* kinetic parameters *V_max_* and *K_m_* at each timestep. The substrate fluxes, together with the lower and upper bounds,*v_l_* and,*v_u_* are used to solve the FBA problem. The FBA problem comprises the stoichiometric matrix *A* and the vector of weights *c*, which represents the contribution of each reaction to the objective function, typically the maximization of the growth rate, *µ*. This optimization is performed iteratively at each timestep *dt* for each species *i*, using the current concentrations of biomass *X_i_*, substrate flux 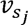 and product flux 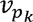 for product *k*.

#### 2.2.1 Calculation of substrate uptake rate

The substrate uptake limit is limited by two factors – the amount of nutrient available in the medium for each species *i* and the transport kinetics for the substrate. We have used a similar The substrate uptake limit is limited by two factors – the amount of nutrient available in the approach as (Chiu, Levy, and Borenstein 2014; Varma and Palsson 1994; Wagner 2022) to *S_j_*(*t*), where the maximum amount of nutrient *j* that species *i* can import per gram biomass per calculate the substrate uptake rate. The nutrient concentration in the medium is denoted by hour is *S_jconc_*

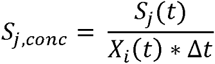

The second limitation is transport, where the cell’s transport mechanism may not be able to import all the available nutrients. Nutrient transport often follows Michaelis–Menten kinetics (Button 1991; Fiksen, Follows, and Aksnes 2013), and we therefore use the kinetic parameters *v_max_* and *K_m_* to calculate the transport limit *S_j,trans_*

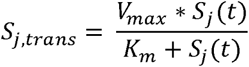

The substrate uptake rate *v_s_* is the minimum of these two limits and is given by

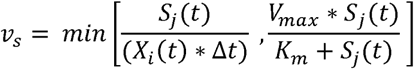

This substrate uptake rate is used as an additional constraint for the parsimonious flux balance analysis to compute the growth rate μ*_i_* of each species *i* and the medium concentration for each substrate is updated.

#### 2.2.2 Flux Variability Analysis (FVA)

To evaluate the production of each product under study we performed an FVA at each time step. The maximum flux through the exchange reaction for *k* products 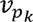 is evaluated as follows

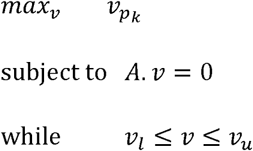

The product concentration *P_k,i_* for product *k* in species, is given by

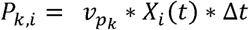

### 2.3 Comparative analysis

High-quality GSMMs of more than 30 organisms were obtained from the BiGG models database (King et al. 2016) and the BioModels database (Malik-Sheriff et al. 2020). These models were filtered based on pathogenicity, model quality and annotation compatibility (Lieven et al. 2020). We chose ten organisms for our final analysis, and though some of these are pathogenic, bioproduction has been successfully demonstrated in all of them. These models organisms, representing the external medium. The uptake kinetic parameters *_Vmax_* and *K_m_* were were used to form pairwise communities with a universal compartment shared by the assumed to be constant (*V_max_* = 20 mmol/gDW/hr and *K_m_* = 0.05 mmol) for all metabolites, as the parameters for all the organisms were not readily available. This value of *V_max_* and *K_m_* lies within the broad range reported in the BRENDA database (Scheer et al. 2011) and has been utilised in several previous studies (Chiu, Levy, and Borenstein 2014; Roman and Wagner 2018). The pairwise community and the monoculture growth rates were simulated in the same external medium, and the product concentrations in both cases were compared to find the best system under each scenario and target product, as shown in Figure 2. The final biomass, substrate, and product concentrations were averaged over the last five feasible solutions to ensure robustness. A minimal abundance of 0.1 and at least a 10% increase in biomass was set as a requirement for the community to be considered viable.

**Fig 2.**
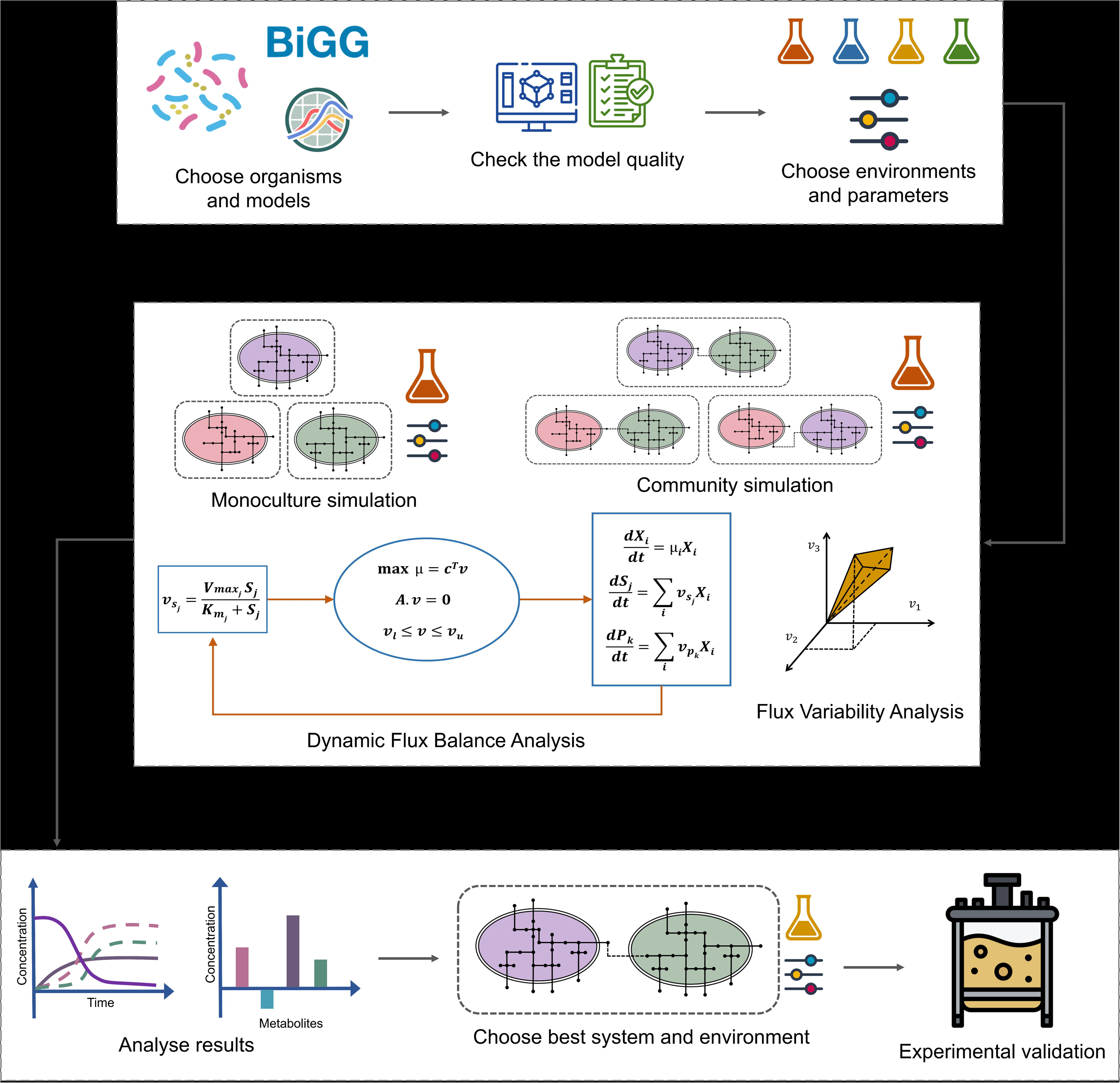
Workflow of COSMOS. The preprocessing step involves selecting target organisms, ensuring model quality, and setting parameters like,, initial biomass, and process time. The algorithm uses these inputs to form pairwise communities and applies dynamic FBA and FVA to simulate growth and productivity across mono- and co-cultures in the same environment. The resulting growth, abundance, and productivity metrics help identify the optimal microbial system for the given conditions, which can then be experimentally validated for bioprocess implementation.

### 2.4 Statistical analysis

The distribution of the data was assessed using the Shapiro-Wilk test, which indicated a non-normal distribution. Therefore, non-parametric statistical methods were applied. We implemented a generalized additive model (GAM) regression in R (mgcv package) to model the relationships between variables before hypothesis testing. This approach allowed us to account for and exclude confounding factors before performing statistical comparisons. For group-wise statistically significant difference was detected (*p* < 0.05), we conducted post-hoc pairwise comparisons, we used the Kruskal-Wallis test, a non-parametric alternative to ANOVA. If a comparisons using the Dunn test with Benjamini-Hochberg correction to adjust for multiple comparisons. All statistical analyses were performed in R (version 4.4.2) using the mgcv and stats packages.

## 3 Results

### 3.1 Navigating Trade-offs: Choosing Between Monocultures and Communities

The medium composition is one of the most substantial factors that can affect a bioprocess. We come across both nutrient-dense or ‘rich’ feedstock like animal manure and sludge, which have more nitrogen content and trace minerals, and ‘less-dense’ feedstock like wheat straw and corn stover (Mohd Johari et al. 2020), which may lack essential micronutrients. The biosynthetic capability of the organisms can also be affected as a result of this medium composition (Williams et al. 2016). So, we need to either choose the target product based on the feedstock availability or source the best-suited feedstock for our product of interest. Using COSMOS, we analyse the effect of medium composition on the biosynthetic capability of a diverse set of organisms.

To investigate the impact of medium composition, we designed two types of media: a ‘minimal medium’ and a ‘rich medium.’ The ‘rich medium’ contains over 60 metabolites commonly found in growth media (Pacheco, Moel, and Segrè 2019). In contrast, the minimal medium consists of around 40 essential metabolites required for the growth of all the individual organisms under study. A complete list of organisms and media components is provided in Supplementary Tables 1-3.

The initial concentration of each component was set to 10 mmol/L in both media. We also investigated the impact of oxygen availability by growing communities under both anaerobic and aerobic conditions, with the latter set to an oxygen concentration of 10 mmol/L. Obligate anaerobes and aerobes were excluded from their respective simulations (Supplementary Table 1), resulting in four distinct environments: aerobic-rich, aerobic-minimal, anaerobic-rich, and anaerobic-minimal. An initial biomass of 0.01 g/L was chosen for both the organisms of the co-culture to ensure sufficient growth. All simulations were run for 12 hours, by which the carbon source was fully consumed, and the organisms reached the stationary phase. We calculated the productivity of each product for both communities and the constituent monocultures, as shown in Figure 3.

**Figure 3.**
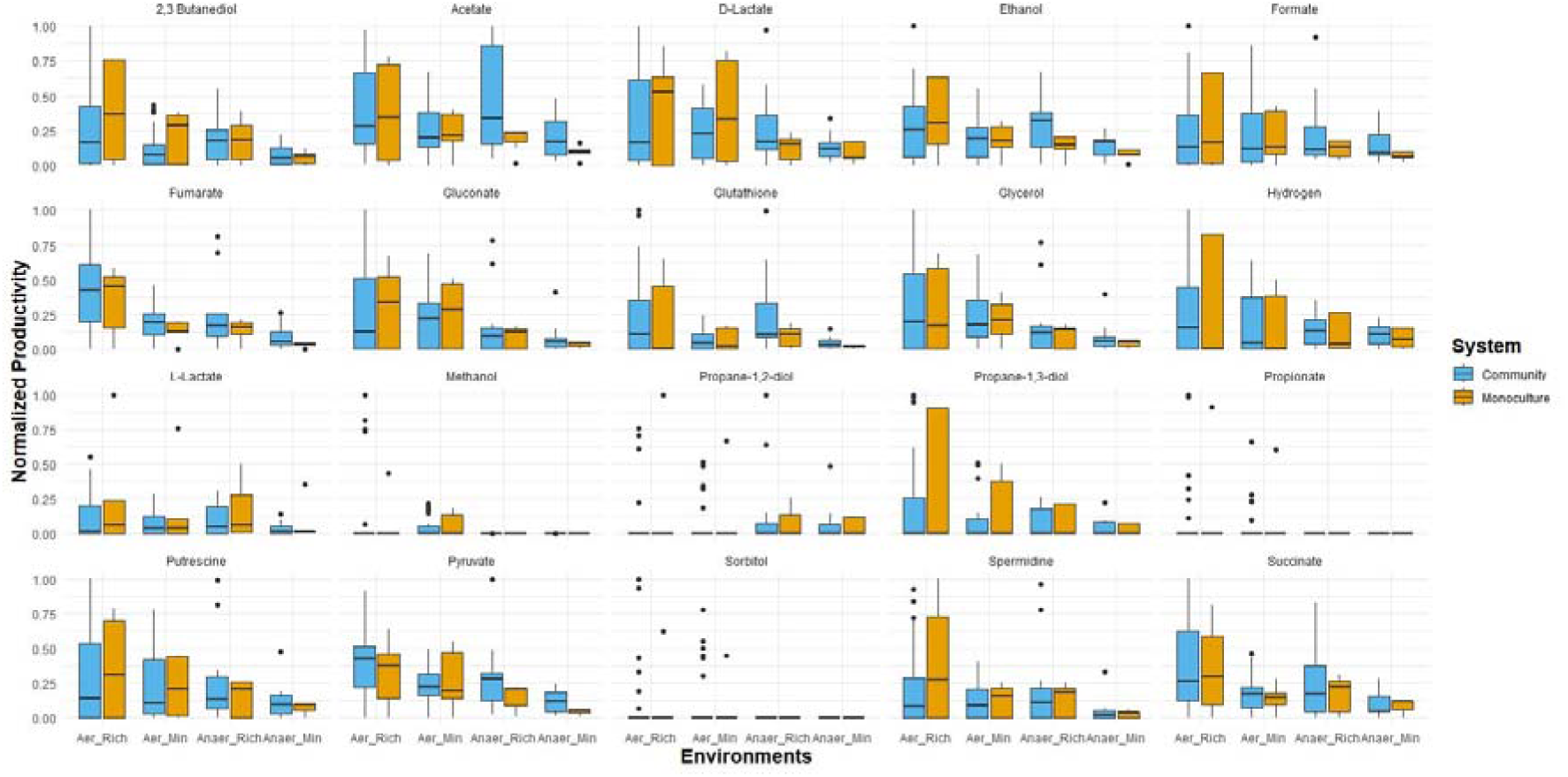
Productivity of communities and monocultures across varying environmental conditions. The productivity of both communities and monocultures for products across all four environments is represented. Five products, viz. adipic acid, xylitol, catechol, butanol and butyrate, were excluded as they are not produced in any of the organisms under study.

When comparing the four environments, we found that overall productivity is highest in the aerobic-rich environment (Figure 3). However, we found the effect of the environment on microbial systems to be more nuanced and product-dependent. While monocultures, on average, performed better in the aerobic-rich environment, communities often achieved the highest productivity for specific products. The list of best-performing microbial systems for the aerobic-rich environment is summarized in Table 1. Our results highlight the importance of conducting an analysis tailored to the specific product and environmental conditions to identify the optimal microbial system.

**Table 1.**
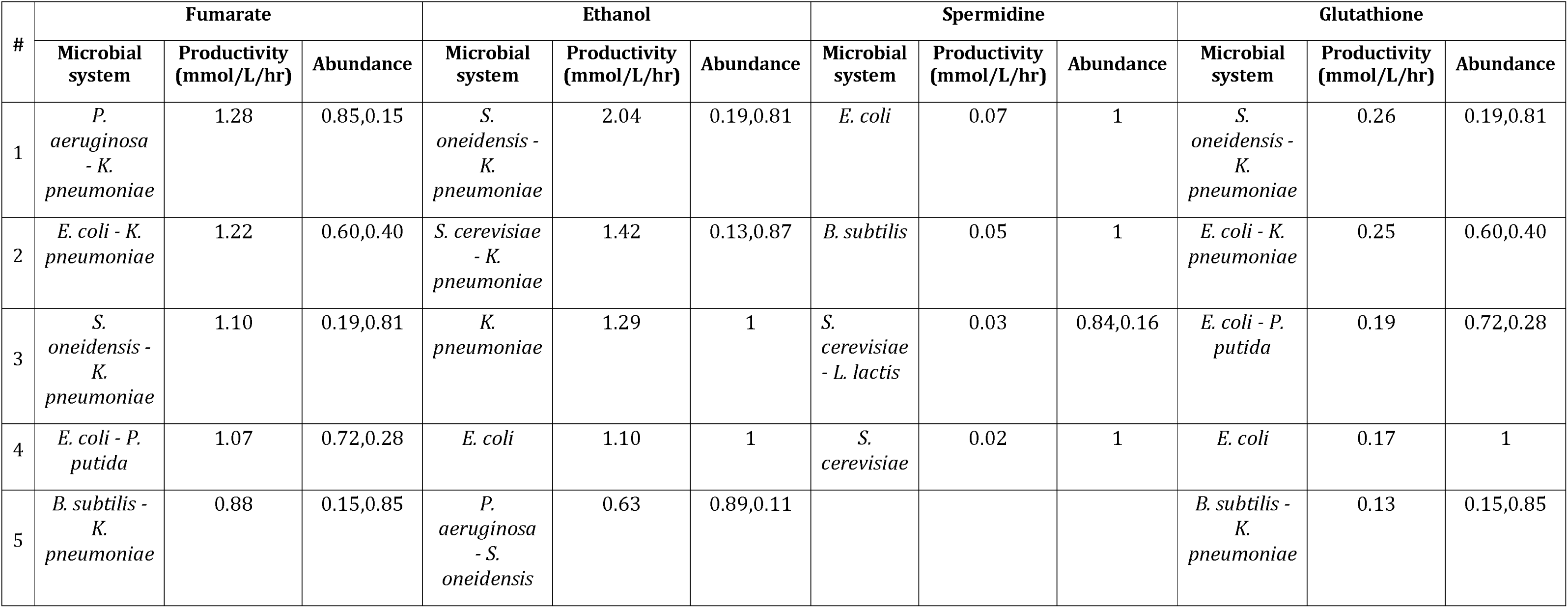
Optimal microbial for four products in the aerobic-rich environment.

| # | Fumarate |  |  | Ethanol |  |  | Spermidine |  |  | Glutathione |  |  |
| --- | --- | --- | --- | --- | --- | --- | --- | --- | --- | --- | --- | --- |
|  | Microbial system | Productivity (mmol/L/hr) | Abundance | Microbial system | Productivity (mmol/L/hr) | Abundance | Microbial system | Productivity (mmol/L/hr) | Abundance | Microbial system | Productivity (mmol/L/hr) | Abundance |
| 1 | *P. aeruginosa* - *K. pneumoniae* | 1.28 | 0.85,0.15 | *S. oneidensis* - *K. pneumoniae* | 2.04 | 0.19,0.81 | *E. coli* | 0.07 | 1 | *S. oneidensis* - *K. pneumoniae* | 0.26 | 0.19,0.81 |
| 2 | *E. coli* - *K. pneumoniae* | 1.22 | 0.60,0.40 | *S. cerevisiae* - *K. pneumoniae* | 1.42 | 0.13,0.87 | *B. subtilis* | 0.05 | 1 | *E. coli* - *K. pneumoniae* | 0.25 | 0.60,0.40 |
| 3 | *S. oneidensis* - *K. pneumoniae* | 1.10 | 0.19,0.81 | *K. pneumoniae* | 1.29 | 1 | *S. cerevisiae* - *L. lactis* | 0.03 | 0.84,0.16 | *E. coli* - *P. putida* | 0.19 | 0.72,0.28 |
| 4 | *E. coli* - *P. putida* | 1.07 | 0.72,0.28 | *E. coli* | 1.10 | 1 | *S. cerevisiae* | 0.02 | 1 | *E. coli* | 0.17 | 1 |
| 5 | *B. subtilis* - *K. pneumoniae* | 0.88 | 0.15,0.85 | *P. aeruginosa* - *S. oneidensis* | 0.63 | 0.89,0.11 |  |  |  | *B. subtilis* - *K. pneumoniae* | 0.13 | 0.15,0.85 |

To efficiently compare the productivity of communities and monocultures, we calculated the productivity ratio for all products across the four environments. It should be noted that the maximum productivity of the two monocultures was compared against the productivity of the community. The productivity ratios across all four environments are represented in Figure 4.

**Fig 4.**
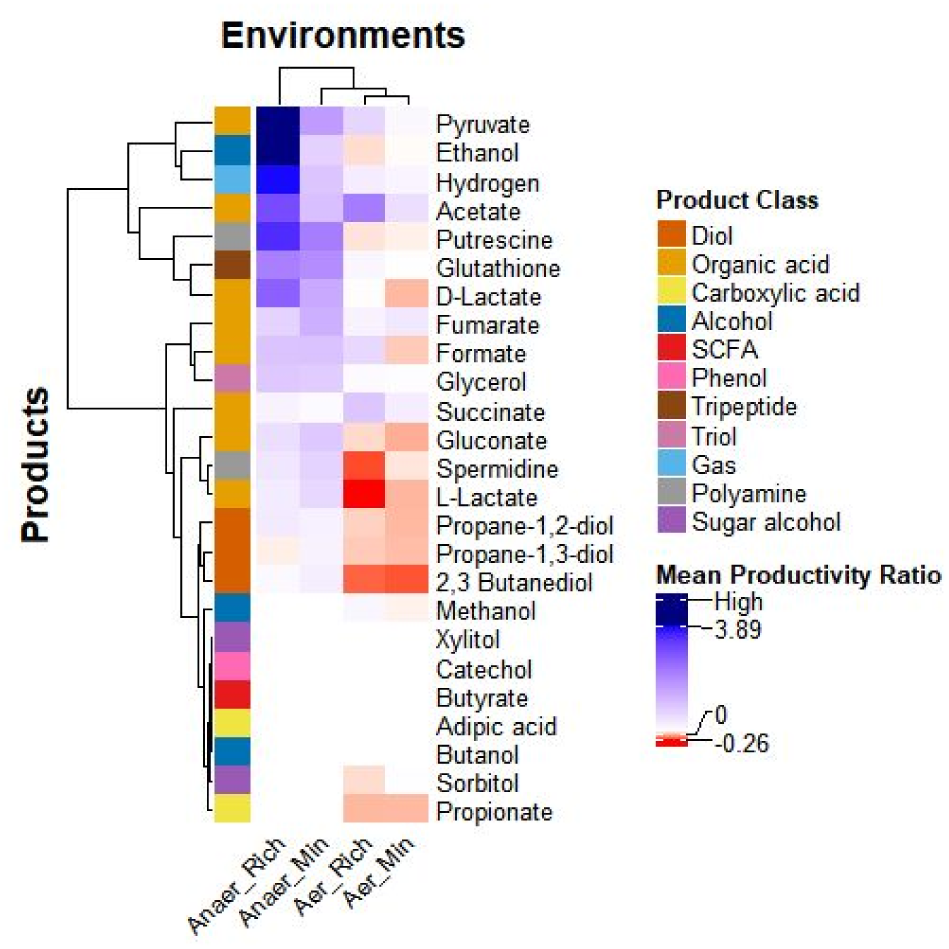
Effect of medium composition and oxygen availability on the productivity of communities vs monocultures. The average productivity ratio of communities across all products and environments is represented. Positive values (blue) indicate increased productivity in communities compared to monocultures, while negative values (red) indicate a decline. In cases where monoculture productivity is extremely low, leading to an exceptionally high productivity ratio, we classify it as ‘High’ and denote it with a navy blue colour.

In a community with organisms A and B, where

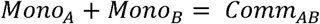

the change in productivity is given as

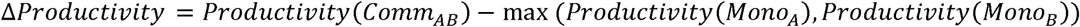

and the ratio of productivity of the community to that of the monoculture is given as

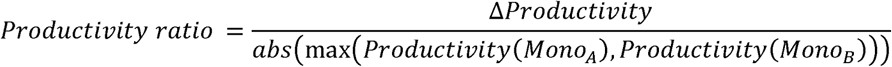

When comparing the productivity ratio of communities and monocultures, we find that the anaerobic-rich environment predominantly favours community-based production, while monocultures perform best in the aerobic-minimal medium (*p*= 2.38×10^-10^; Supplementary Data 5). One possible explanation is that the incomplete nature of anaerobic fermentation can lead to lower productivity in monocultures. In contrast, microbial communities may exchange primary metabolites, enhancing both growth and productivity, particularly in anaerobic conditions. To test this hypothesis, we analysed the abundance ratios of five communities across all four environmental conditions, as shown in Table 2. Under anaerobic conditions, although total biomass decreases, the abundance ratios of both organisms become more balanced. This equilibrium may contribute to the improved performance of communities in anaerobic environments. Additionally, the rich medium supports higher productivity by promoting better growth and increased metabolite exchange. Based on these findings, simple monocultures may be preferable for aerobic environments, while communities offer advantages for anaerobic fermentation.

**Table 2.** Comparison of Microbial Communities Exhibiting Growth Across All Environments.

|  | Aerobic-rich |  |  | Aerobic-minimal |  |  | Anaerobic-rich |  |  | Anaerobic-minimal |  |  |
| --- | --- | --- | --- | --- | --- | --- | --- | --- | --- | --- | --- | --- |
| Community | Biomass | Total Biomass | Abundance | Biomass | Total Biomass | Abundance | Biomass | Total Biomass | Abundance | Biomass | Total Biomass | Abundance |
| *E. coli* - *L. lactis* | 0.76,0.17 | 0.93 | 0.82,0.18 | 0.57,0.13 | 0.7 | 0.81,0.19 | 0.35,0.23 | 0.58 | 0.60,0.40 | 0.19,0.24 | 0.43 | 0.44,0.56 |
| *L. lactis* - *S. oneidensis* | 0.13,0.35 | 0.48 | 0.26,0.74 | 0.13,0.36 | 0.49 | 0.26,0.74 | 0.14,0.22 | 0.36 | 0.38,0.62 | 0.13,0.22 | 0.35 | 0.38,0.62 |
| *L. lactis* - *K. pneumoniae* | 0.12,0.80 | 0.92 | 0.13,0.87 | 0.12,0.35 | 0.47 | 0.15,0.85 | 0.13,0.48 | 0.61 | 0.21,0.79 | 0.14,0.29 | 0.43 | 0.32,0.68 |
| *B. subtilis* - *K. pneumoniae* | 0.14,0.80 | 0.94 | 0.15,0.85 | 0.14,0.66 | 0.8 | 0.18,0.82 | 0.17,0.46 | 0.63 | 0.27,0.73 | 0.17,0.29 | 0.46 | 0.37,0.63 |
| *S. oneidensis* - *K. pneumoniae* | 0.19,0.78 | 0.97 | 0.19,0.80 | 0.15,0.64 | 0.79 | 0.19,0.81 | 0.23,0.48 | 0.71 | 0.32,0.68 | 0.25,0.27 | 0.52 | 0.47,0.53 |

However, as discussed earlier, there are notable exceptions where communities excel in aerobic environments. This underscores the importance of evaluating all candidate organisms as monocultures and communities to determine the optimal microbial system for a given product and environment. If the differences in productivity are minimal, monocultures may be preferable due to easier process control, whereas co-cultures can be advantageous if they offer superior biosynthetic capabilities.

For example, in the aerobic-rich medium, the sorbitol productivity of the *S. cerevisae – B. subtilis* and the *S . c e r e v i s a e –* c*L*om. m*l* u*a*ni*c*ti*t*es*i s*are the highest and have 59% and 48% improvement, respectively, when compared to monocultures. Likewise, in the anaerobic-minimal medium*, Bacillus* subtilis outperforms all the other microbial communities in the production of L-lactate, with the productivity of B. subtilis being 0.29 mmol/L/hr. It is to be noted that the second-best alternative is *E. coli – P. aeruginosa* community, with a productivity of 0.12 mmol/L/hr of L-lactate, which is less than half of the monoculture. These examples demonstrate that while general trends provide useful insights, exceptions can be strategically utilized to optimize bioprocesses.

Therefore, a systematic screening of both monocultures and communities is essential. In the following sections, we explore the impact of additional factors, such as interaction type, carbon sources and initial biomass ratio, on productivity and community dynamics.

### 3.2 Synergy in Action: Positive Interactions Drive Higher Productivity

As we are dealing with pairwise communities, another interesting factor that we can analyse is the effect of interaction. To find the interaction type of a community, we compare the growth of the organism as a monoculture and in the community under the same environmental conditions. The communities were categorised into six interaction types, namely, competition, amensalism, parasitism, neutralism, commensalism and mutualism, based on a 10% difference in growth rates of the microbe in the co-culture and the monoculture (Heinken and Thiele 2015).

The average Productivity ratio was calculated for each product across the four environments, is shown in Figure 5. Mutualistic interactions generally lead to the highest productivity gains in communities, likely due to the increased biomass of the participating organisms. Parasitic interactions follow, with productivity varying depending on the product— some are better produced in communities, while others are more efficiently generated in monocultures. This variation may arise because, in some cases, the producer strain exhibits higher growth, whereas in others, it experiences reduced growth. The effect of interaction type on the productivity ratio was statistically significant (p = 1.4 x 10^-10^; Supplementary Data 6).

**Fig 5.**
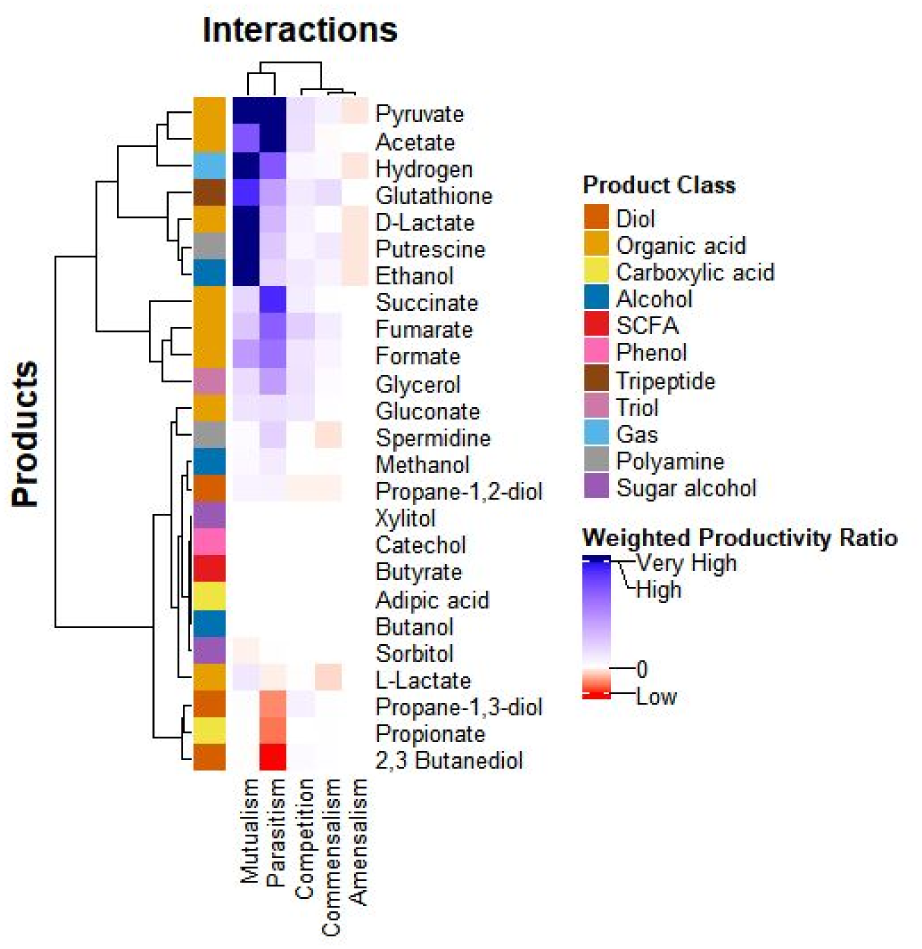
Effect of interaction type on the productivity of communities vs monocultures across products. Positive values (blue) indicate an increase in the productivity of a given product in the community compared to the monocultures, while negative values (red) reflect a decline in productivity. Neutralism was not observed.

However, no significant differences were observed among competition, commensalism, and amensalism, suggesting that these interaction types have minimal impact on productivity differences between communities and monocultures.

### 3.3 Tailored Metabolism: Communities and Monocultures Show Distinct Carbon Preferences

Another key factor we examined is the impact of different carbon sources on the growth and biosynthetic capabilities of microbial systems. In this analysis, we replaced the glucose in the ‘rich’ and ‘minimal’ media with 10 mmol/L of the carbon source under investigation. We tested seven different carbon sources and compared the productivity of communities and monocultures across all four environments, as shown in Figure 6.

**Fig 6.**
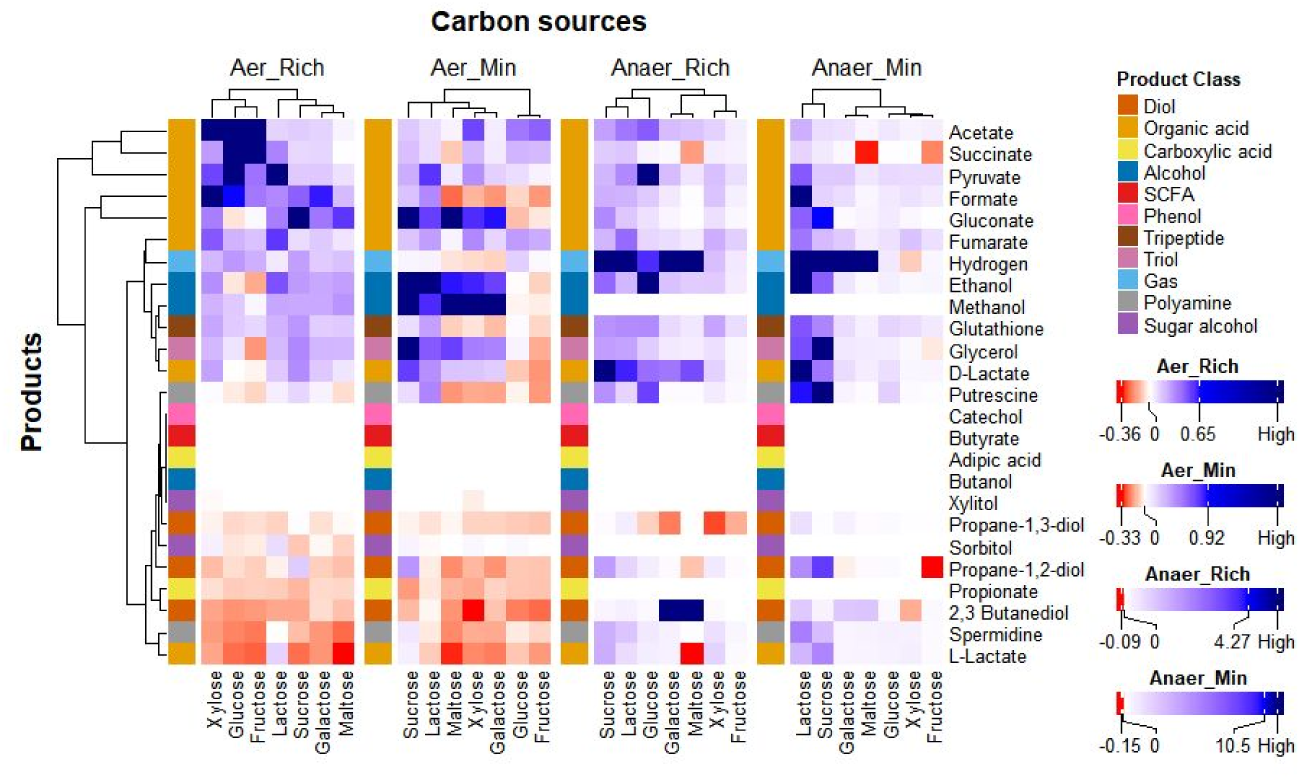
Effect of carbon source on the productivity of communities vs monocultures. The effect of carbon sources on the Productivity ratio is compared across all four environments. Positive values (blue) indicate an increase in the productivity of a given product in the community compared to the monocultures, while negative values (red) reflect a decline in productivity.

Our findings show that while there are some variations, lactose and sucrose consistently enhance productivity in communities, whereas xylose and fructose favour monocultures. Interestingly, the aerobic-rich medium deviates from the pattern observed in the other three environments. While xylose and fructose promote higher productivity in communities in the aerobic-rich environment, these carbon sources support higher productivity in monocultures in other environments. Although the underlying cause remains unclear, this provides an additional factor that could be manipulated to optimize the bioprocess. This insight can guide the design or supplementation of fermentation media such that the choice of carbon source aligns with the microbial system and environment. The effect of carbon source on the productivity ratio was statistically significant (p = 1.8 x 10^-4^; Supplementary Data 7).

To explore the variation amongst communities, we calculated the productivity ratio for four different products across communities in the aerobic-rich medium, as shown in Supplementary Figure 1. Some communities, like *Shewanella oneidensis – Klebsiella pneumoniae*, showed minimal sensitivity to various carbon sources. However, others, such as *S. cerevisiae – L. lactis,* displayed varying productivity across carbon sources. In some cases, like *E. coli – S. cerevisiae*, the community does not grow under specific carbon sources. Moreover, this behaviour can also be product-dependent. Thus, product-specific analysis is crucial for identifying the optimal combination of carbon source and microbial system.

### 3.4 Computational Screening Reveals Ideal Microbial Systems Across Fermentation Conditions

While communities generally perform better in challenging environments, exceptions may occur. Therefore, it is essential to analyse the specific product of interest before selecting the most suitable microbial system. Furthermore, variations between communities mean that choosing the optimal community for a given nutrient source is crucial. Since experimentally comparing multiple communities is time-consuming and labour-intensive, computational analysis becomes essential.

We compared the change in productivity between communities and monocultures across all products, as shown in Figure 8. The mean productivity ratio for 25 products was calculated across the four environments for each community. Positive values (blue) indicate higher productivity in communities compared to their constituent monocultures, while negative values (red) indicate lower productivity. Clear clustering patterns emerge both by product and community, highlighting preferences in microbial systems. The effect of community on the productivity ratio was found to be statistically significant, with a (p = 2.2 x 10^-16^;Supplementary Data 8).

**Fig 7.**
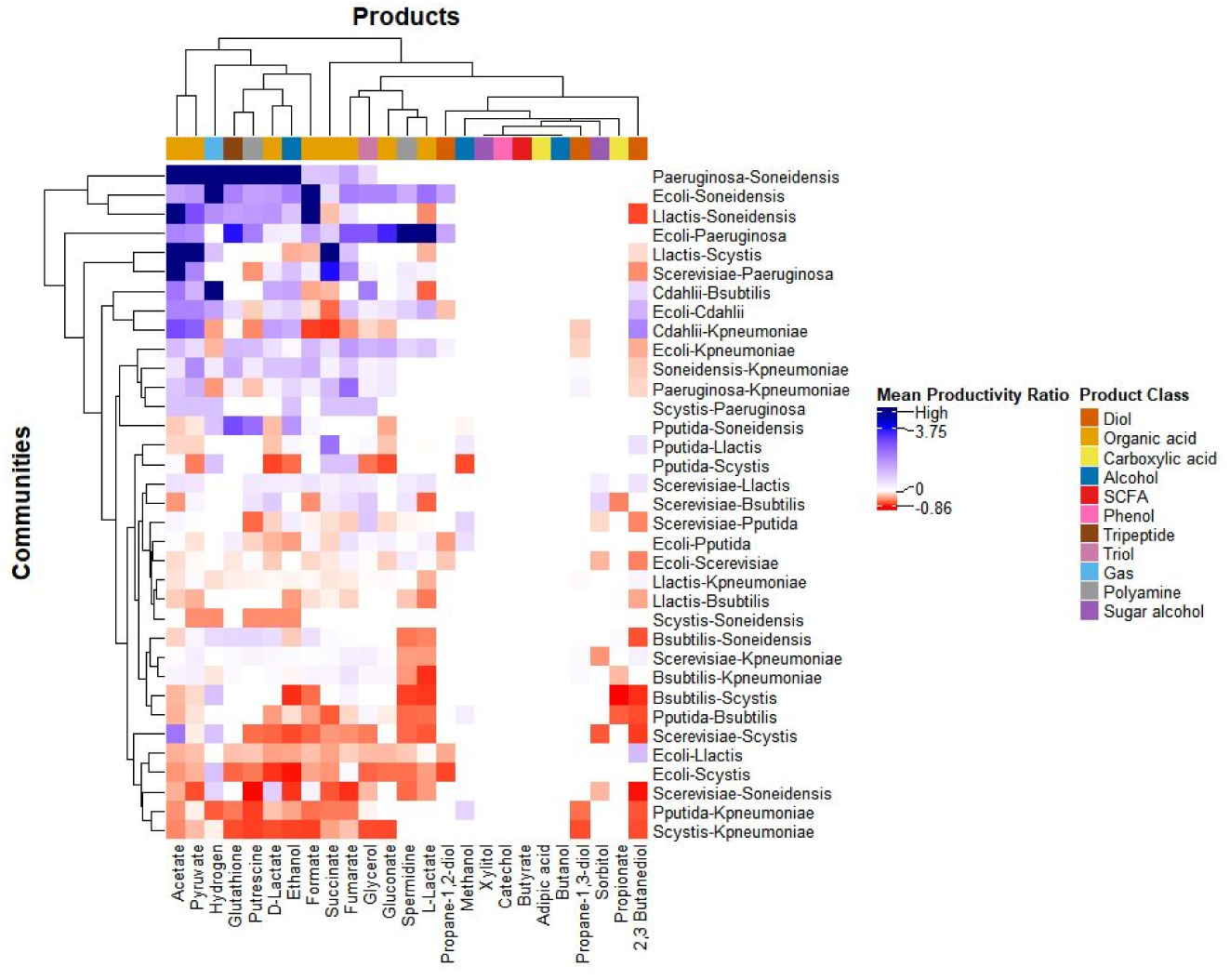
Comparison of Productivity of Communities and Monocultures. This figure presents the mean productivity ratio of communities, averaged across four environments, for all the products under study. Positive values (blue) indicate an increase in the productivity of a given product in the community compared to the monocultures, while negative values (red) reflect a decline in productivity.

Microbial communities such as *P. aeruginosa–S. oneidensis* and *E. coli–S. oneidensis* generally outperform their monocultures, particularly in the production of metabolites like pyruvate, hydrogen, and glutathione. In contrast, communities like *Synechocystis spp.–K. pneumoniae* and *P. putida–K. pneumoniae* consistently exhibit lower productivity compared to their respective monocultures. Furthermore, metabolite production appears to be highly product- and organism-dependent. For instance, acetate and pyruvate are more efficiently synthesized in co-culture systems, whereas sorbitol and propionate yield higher levels in monocultures. Additionally, productivity patterns vary across different environmental conditions, underscoring the importance of evaluating microbial systems in a context-specific manner.

To investigate this behaviour further, we identified the best microbial systems for producing four key products from different classes under all four environmental conditions, as summarized in Supplementary Table 5. The observations reveal that productivity is highest in the aerobic-rich environment, as discussed in Sections 3.1 and 3.2. Notably, the anaerobic-rich environment also performs competitively and, in some cases, surpasses the aerobic minimal medium in productivity. The findings in Supplementary Table 5 show that some of the best-performing systems in aerobic environments are monocultures, while communities tend to outcompete monocultures in anaerobic environments. It also highlights variability within products—while fumarate is consistently better produced by communities across all environments, spermidine is predominantly produced by monocultures in most conditions. This highlights the necessity of selecting the optimal microbial consortium and environmental setting tailored to the specific product of interest.

Moreover, when choosing a community, the selection criteria need not be restricted to productivity but can also be extended to other factors like abundance ratios. When two systems exhibit similar productivity, choosing the one with a more balanced abundance may be preferable, as it can lead to a more stable community. Moreover, the final choice can also be guided by biological insights, such as ease of handling or specific biological characteristics desirable for the bioprocess of interest. The optimal systems in the other environments are provided in Supplementary Data 1-4.

Additionally, we compiled the top-performing systems for all 25 products across four environments in Supplementary Table 6. As expected from previous analyses, monocultures dominate in the aerobic-rich medium. Meanwhile, communities excel in the anaerobic-minimal medium, reinforcing the idea that harsher conditions favour biosynthesis in the communities. The detailed results for all 25 products are in Supplementary Data 1-4, with key findings aligning with experimental studies summarized in Table 3 alongside references.

**Table 3.** Results that align closely with experimental findings.

| # | Product | System Name | Productivity in Community (mmol/L/hr) | Productivity in Monoculture A (mmol/L/hr) | Productivity in Monoculture B (mmol/L/hr) | Improvement in productivity (%) | Abundance | Environment | Reference |
| --- | --- | --- | --- | --- | --- | --- | --- | --- | --- |
| 1 | 1,3 - Propanediol | *S. oneidensis* - *K. pneumoniae* | 0.064 | 0 | 0.0519 | 23.31 | 0.32,0.68 | Anaerobic-rich | (Wang et al. 2023) |
| 2 |  | *P. aeruginosa* - *K. pneumoniae* | 0.053 | 0 | 0.018 | 196.11 | 0.58,0.42 | Anaerobic-minimal | (Li, Zhu, and Ge 2019) |
| 3 | 2,3 - Butanediol | *P. aeruginosa* - *K. pneumoniae* | 0.073 | 0 | 0.036 | 103.06 | 0.58,0.42 | Anaerobic-minimal | (Li, Zhu, and Ge 2019) |
| 4 | Hydrogen | *C. ljungdahlii* - *B. subtilis* | 0.684 | 0.091 | 0 | 647.92 | 0.87,0.13 | Anaerobic-rich | (Chang et al. 2008) |
| 5 | Ethanol | *S. cerevisiae* – *B. subtilis* | 0.630 | 0.416 | 0.604 | 4.41 | 0.69,0.31 | Aerobic-rich | (Tantipaibulvut et al. 2015) |
|  |  | *S. cerevisiae* - *L. lactis* | 0.584 | 0.416 | 0.009 | 40.22 | 0.84,0.16 | Aerobic-rich | (Carvalho et al. 2015) |
|  |  | *S. cerevisiae* - *L. lactis* | 0.410 | 0.363 | 0.008 | 13.03 | 0.65,0.35 | Aerobic-minimal | (Li, Zhu, and Ge 2019) |

### 3.5 Optimising Biomass Ratios Enhances Productivity in Microbial Communities

The initial biomass ratio of organisms within a community can significantly influence growth, abundance, and overall community dynamics. By adjusting the inoculum ratio, we can further enhance the biosynthetic capabilities of the community. Tables 4 and 5 illustrate how variations in inoculum ratios impact 1,3-propanediol (1,3-PDO) production in *P. aeruginosa - K. pneumoniae* and *S. oneidensis - K. pneumoniae* communities. Product concentrations vary with inoculum ratios, highlighting distinct responses between the two communities. While increasing *K. pneumoniae* inoculum enhances product titer in both, the *S. oneidensis - K. pneumoniae* community shows an increase in total biomass, whereas *P. aeruginosa - K. pneumoniae* displays a growth reduction. Moreover, higher *K. pneumoniae* inoculum in *S. oneidensis - K. pneumoniae* leads to a more skewed abundance ratio. These findings suggest that inoculum ratios can have varied effects on community dynamics and biosynthetic output, emphasizing the need for careful ratio optimization.

**Table 4.** Effect of inoculum ratio on the productivity of 1,3-PDO in *P. aeruginosa - K. pneumoniae* community.

| Biomass ratio | Biomass A (g/L) | Biomass B (g/L) | Total Biomass (g/L) | Abundance | Product concentration (mmol/L) |
| --- | --- | --- | --- | --- | --- |
| 0.9,0.1 | 0.67 | 0.03 | 0.70 | 0.96,0.04 | 0.01 |
| 0.8,0.2 | 0.63 | 0.06 | 0.69 | 0.91,0.09 | 0.02 |
| 0.7,0.3 | 0.59 | 0.09 | 0.68 | 0.87,0.13 | 0.03 |
| 0.6,0.4 | 0.54 | 0.12 | 0.66 | 0.81,0.19 | 0.03 |
| 0.5,0.5 | 0.48 | 0.16 | 0.64 | 0.75,0.25 | 0.04 |
| 0.4,0.6 | 0.39 | 0.19 | 0.58 | 0.67,0.33 | 0.06 |
| 0.3,0.7 | 0.33 | 0.23 | 0.56 | 0.59,0.41 | 0.07 |
| 0.2,0.8 | 0.23 | 0.27 | 0.50 | 0.47,0.53 | 0.08 |
| 0.1,0.9 | 0.14 | 0.32 | 0.46 | 0.31,0.69 | 0.09 |

**Table 5.** Effect of inoculum ratio on the productivity of 1,3-PDO in *S. oneidensis - K. pneumoniae* community.

| **Biomass ratio** | **Biomass A (g/L)** | **Biomass B (g/L)** | **Total biomass (g/L)** | **Abundance** | **Product concentration (mmol/L)** |
| --- | --- | --- | --- | --- | --- |
| 0.1,0.9 | 0.47 | 0.12 | 0.59 | 0.79,0.21 | 0.04 |
| 0.2,0.8 | 0.40 | 0.23 | 0.62 | 0.64,0.36 | 0.07 |
| 0.3,0.7 | 0.35 | 0.33 | 0.68 | 0.51,0.49 | 0.09 |
| 0.4,0.6 | 0.29 | 0.41 | 0.7 | 0.41,0.59 | 0.11 |
| 0.5,0.5 | 0.23 | 0.48 | 0.71 | 0.32,0.68 | 0.14 |
| 0.6,0.4 | 0.19 | 0.57 | 0.76 | 0.24,0.76 | 0.14 |
| 0.7,0.3 | 0.13 | 0.63 | 0.76 | 0.18,0.82 | 0.18 |
| 0.2,0.8 | 0.09 | 0.67 | 0.76 | 0.11,0.88 | 0.22 |
| 0.9,0.1 | 0.04 | 0.75 | 0.8 | 0.05,0.95 | 0.23 |

### 3.6 Validation

Though some of the results we obtain corroborate with experimental studies as listed in Table 3, these studies only talk about the biosynthetic capability of the community and not the monoculture. There are very few studies that compare the production capabilities of the monoculture and the community in the same environment. In one such study (Wang et al. 2023), they evaluated the production of 1,3-PDO in *Klebsiella pneumoniae* monoculture and the *Klebsiella pneumoniae – Shewanella oneidensis* community using glycerol as the carbon source under anaerobic conditions. They found that *Shewanella oneidensis* acts as an electron mediator in the co-culture to attain better production of 1,3-PDO. Notably, under anaerobic conditions, this community is the top producer of 1,3-PDO in our analysis. To demonstrate the reliability of COSMOS, we simulated the growth of the community and the monoculture in the medium used in the experimental study and compared the results. The first analysis is to identify the optimum glycerol concentration for the monoculture. We found 50 g/L glycerol to be the optimum concentration for 1,3-PDO production, which correlates with the experiment values, as shown in Table 6.

**Table 6.** Concentration of 1,3 propanediol in experiments and simulations under varying glycerol concentration.

| Glycerol concentration (g/L) | Concentration of 1,3 PDO in experiments (g/L) | Concentration of 1,3 PDO in COSMOS (g/L) |
| --- | --- | --- |
| 40 | 11.3 | 1.76 |
| 50 | 12.68 | 1.83 |
| 70 | 12.21 | 1.58 |

For the community analysis, the experimental group used a fed-batch reactor under anaerobic conditions where the initial glycerol concentration was 50 g/L. When the glycerol concentration falls below 10 g/L, additional glycerol is introduced to maintain a final concentration of 30 g/L. They found that the community produced 32.01 g/L of 1,3-PDO, which was significantly higher when compared to the monoculture. They also optimised the inoculum ratio and found that a 1:1 ratio of *Klebsiella pneumoniae* : *Shewanella oneidensis* resulted in the highest product concentration, though the exact values are not listed. We did a similar analysis using COSMOS and found that the community was, in fact, able to produce more product in the fed-batch system when compared to the monoculture. However, we found that both the inoculum ratios of 1:1 and 2:1 performed equally well, as shown in Table 7. This might be because we have used the wild-type models of both organisms and lack the strain-specific information that might be required.

**Table 7.**
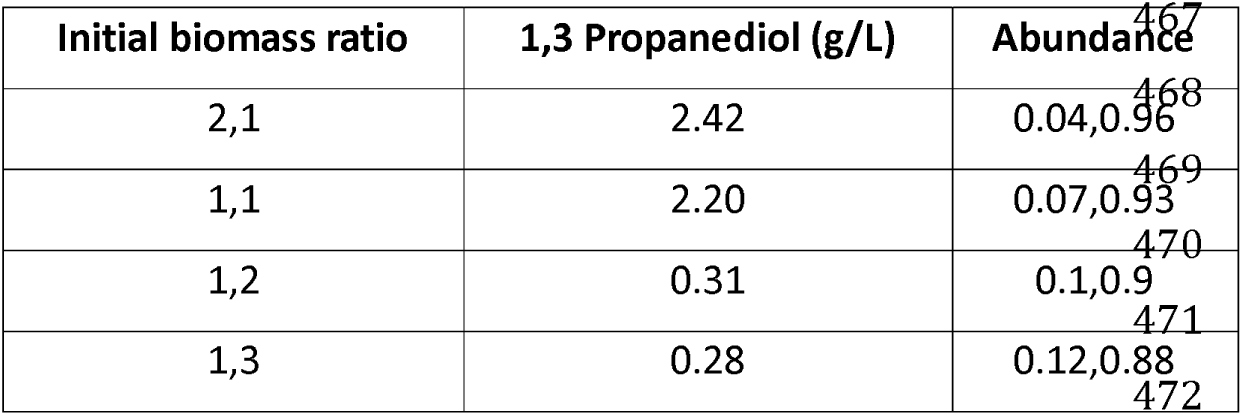
Concentration of 1,3 -propanediol with varying inoculum ratio according to simulations.

| Initial biomass ratio | 1,3 Propanediol (g/L) | Abundance |
| --- | --- | --- |
| 2,1 | 2.42 | 0.04,0.96 |
| 1,1 | 2.20 | 0.07,0.93 |
| 1,2 | 0.31 | 0.1,0.9 |
| 1,3 | 0.28 | 0.12,0.88 |

The study reported monoculture and co-culture yields of 0.4 g/g and 0.44 g/g, respectively, reflecting a 10% improvement. COSMOS predicted yields of 1.50 g/g for monocultures and 1.33 g/g for co-cultures, indicating a 12.6% increase. Although the absolute values differ, the algorithm effectively captured the trend in performance, identifying optimal substrate concentrations and inoculum ratios. This demonstrates the potential of dynamic modelling in bioprocess design, offering a powerful tool for efficiently comparing multiple communities and monocultures to identify the most productive configurations.

## 4 Discussion

Microbial communities often exhibit superior biosynthetic capability compared to monocultures (Rapp, Jenkins, and Betenbaugh 2020; Xu and Tschirner 2011). However, their application in bioproduction remains limited to cases where their productivity is already known or where their metabolic diversity is essential, such as when one community member can metabolise a complex carbon source that the others cannot. Studies that directly compare the biosynthetic potential of multiple communities remain scarce (Wilken et al. 2018), leaving a gap in identifying the most effective microbial systems.

To address this, COSMOS provides a systematic approach to evaluating microbial systems by assessing a wide range of co-cultures alongside their corresponding monocultures. Unlike tools like FLYCOP (García-Jiménez, García, and Nogales 2018), which optimizes a single co-culture, COSMOS evaluates a broad range of co-cultures alongside their constituent monocultures to identify the most effective microbial system for a given product. It also offers granular insights into the system’s performance across a wider range of environmental conditions. COSMOS is particularly valuable when working with low-quality feedstocks, such as agricultural residues and wastewater, which are gaining importance due to global efforts to transition toward second-generation feedstocks for improved food security and sustainability (Russo et al. 2025). Given these challenges, optimizing the microbial system is just as crucial as optimizing the environment to develop efficient and sustainable bioprocesses.

While productivity is often the primary factor in selecting a microbial system, the choice between monocultures and communities depends on additional factors. If a community offers substantially higher productivity, it is the better option for bioproduction. However, when productivity is comparable, monocultures are preferred for their ease of control and simplicity. In cases where operational simplicity takes precedence over maximum output, slightly less productive monocultures can be selected. COSMOS streamlines this decision-making process by assessing the biosynthetic potential of diverse microbial systems under diverse environments.

COSMOS employs parsimonious dynamic FBA (Henson and Hanly 2014; Mahadevan, Edwards, and Francis J Doyle 2002) to model microbial growth in synthetic consortia, allowing it to capture fluctuating growth rates. While traditional steady-state modelling is simpler to use, it assumes equal growth rates and is better suited to natural communities (Chan, Simons, and Maranas 2017). Therefore, we use dynamic FBA, which accounts for fluctuating growth and provides promising results even with the lack of organism-specific parameters (Gomez, Höffner, and Barton 2014). This makes COSMOS especially valuable for engineered or synthetic consortia, where kinetic data is often unavailable, and the stability of the co-culture remains uncertain.

By integrating Flux Variability Analysis (FVA), COSMOS minimizes redundancy in metabolic predictions, providing more precise insights into productivity (Gudmundsson and Thiele 2010). To ensure an unbiased comparison, COSMOS evaluates communities and monocultures under identical conditions, benchmarking community productivity against the highest-performing monoculture. This approach prioritizes final productivity or yield, making it a more effective tool for selecting optimal bioproduction systems (Zaramela et al. 2021).

To understand the factors that influence bioproduction in cocultures and monocultures, we tested COSMOS across four environments: aerobic-rich, aerobic-minimal, anaerobic-rich, and anaerobic-minimal media. While overall productivity was highest in the aerobic nutrient-rich conditions, communities outperformed monocultures in anaerobic environments. This advantage is likely driven by cooperative interactions facilitated by metabolite exchange from incomplete anaerobic fermentation. Additionally, the more balanced abundance ratios observed in co-cultures under anaerobic conditions further support this hypothesis.

Interestingly, there were exceptions where communities achieved maximum productivity in aerobic environments, while certain monocultures outperformed communities in anaerobic conditions. This highlights the importance of systematically evaluating both communities and monocultures before selecting the optimal microbial system for a specific product. COSMOS enables us to maximize productivity in the chosen fermentation medium and provides insights into the role of mutualistic interactions in community design. By leveraging these interactions, COSMOS aids in optimizing both productivity and stability for biomanufacturing applications.

We also examined the effect of carbon sources on these systems. Certain carbon sources, like lactose and sucrose, enable better productivity in the community, while fructose boosts monoculture performance in most environments. This analysis aids in selecting or enriching fermentation media with specific carbon sources. The algorithm can also be easily extended to study the effect of two or more carbon sources. COSMOS also equips us to optimise the initial inoculum ratio to further improve productivity.

To validate COSMOS, we applied it to the *Klebsiella pneumoniae – Shewanella oneidensis* community and its corresponding monocultures, successfully capturing the effects of carbon source concentration and inoculum ratio (Wang et al. 2023). The observed trends in yield and productivity across different conditions aligned with experimental data, confirming the reliability of the algorithm. Furthermore, several microbial communities identified through our analysis have been experimentally validated as stable consortia, with some already proving their biosynthetic potential (as listed in Table 3).

While COSMOS provides valuable results using standard kinetic parameters and wild-type GSMMs, its accuracy can be significantly improved by using experimentally-determined kinetic parameters and strain-specific GSMMs (C. J. Foster et al. 2021; Link, Christodoulou, and Sauer 2014). However, the availability of high-quality, manually curated models for diverse organisms remains a limitation (Ponce-de-Leon et al. 2015). Improving the accessibility and accuracy of metabolic models will strengthen computational predictions. Additionally, integrating multi-omic data—such as transcriptomic and proteomic information—into GSMMs could further refine the algorithm’s performance, paving the way for more precise bioprocess optimization (Sen and Orešič 2023).

Notably, the insights from this study have direct relevance to fields like biofuel production, wastewater treatment, and pharmaceutical biosynthesis (Keasling et al. 2021). Moreover, by optimising microbial systems for nutrient-limited and waste-derived feedstocks, our approach aligns with the UN Sustainable Development Goals (SDGs), particularly SDG 9 (Industry, Innovation, and Infrastructure) and SDG 12 (Responsible Consumption and Production) (Keasling et al. 2021). In summary, COSMOS provides a robust computational framework to evaluate and compare the productivity of monocultures and microbial communities, addressing the challenges of experimental testing. It offers insights into how environmental factors, carbon sources, and inoculum ratios affect performance, helping researchers make well-informed decisions about which microbial system to use. Whether the goal is to work with nutrient-dense or nutrient-limited media, our approach ensures the selection of the most effective system, enhancing productivity, process efficiency, and community stability.

## 5 Conclusion

As industries pivot toward sustainable practices, bioresources like agricultural residues and wastewater will play a vital role in producing specialty and bulk chemicals. Although nutrient-rich media remain standard for high-value fermentation, optimizing bioprocesses with nutrient-poor media is increasingly crucial for achieving net-zero carbon goals. However, the limited economic feasibility and lack of robust tools have hindered their widespread adoption.

Our study demonstrates that *in silico* analyses can effectively optimise microbial systems to enhance productivity, especially in challenging environments. Microbial communities are generally better suited for anaerobic conditions due to their complementary metabolism and cooperative interactions. However, certain communities can outperform monocultures even in aerobic environments, demonstrating their enhanced biosynthetic capabilities. Likewise, some monocultures may still surpass communities in anaerobic settings, underscoring the importance of systematic computational screening to identify the most suitable candidates before laboratory validation. Tools like COSMOS facilitate this process by helping us leverage both productivity and ease of control, enabling informed decision-making. As more models and kinetic data become available, the predictive accuracy of computational approaches will continue to improve, further advancing bioprocess design.

Ultimately, this approach supports the rational design of microbial systems, advancing synthetic biology and sustainable biomanufacturing. By understanding when to switch between monocultures and communities and identifying key environmental and metabolic factors influencing performance, we can develop more robust and efficient microbial systems. We believe that our framework and such computational strategies will be key enablers in the transition to circular bio-economies.

## Acknowledgements

LR acknowledges the fellowship from the Ministry of Education, Government of India. KR acknowledges support from the Science and Engineering Research Board (SERB) MATRICS Grant MTR/2020/000490, IIT Madras, Centre for Integrative Biology and Systems mEdicine (IBSE) and the Wadhwani School of Data Science and AI.

## Data Availability

All models and analysis scripts used in this study are openly available at https://github.com/RamanLab/COSMOS

## Author Contributions

Lavanya Raajaraam: Conceptualization, Performing Experiments, Analysis of Results, Writing - original draft. Karthik Raman: Conceptualisation, Analysis of Results.

## Declaration of Competing Interest

The authors declare that they have no known competing financial interests or personal relationships that could have appeared to influence the work reported in this paper.

## Notes

http://bigg.ucsd.edu/

https://www.ebi.ac.uk/biomodels/

### Competing Interest Statement

The authors have declared no competing interest.

### Summary of Updates

Visualization has been improved and statistical tests were performed.

