## Supplementary Tables 1-3 for "COSMOS: COmmunity and Single Microbe Optimisation System": Description of Additional Supplementary Data.docx

**Description of Additional Supplementary Files**

Supplementary Data 1 – Aerobic - Rich environment analysis

Supplementary Data 2 – Aerobic - Minimal environment analysis

Supplementary Data 3 – Anaerobic - Rich environment analysis

Supplementary Data 4 – Anaerobic - Minimal environment analysis

Supplementary Data 5 – Statistical analysis of the effect of environment on productivity ratio

Supplementary Data 6 – Statistical analysis of the effect of interaction on productivity ratio

Supplementary Data 7 – Statistical analysis of the effect of carbon source on productivity ratio

Supplementary Data 8 – Statistical analysis of the effect of community on productivity ratio
