## Supplementary Tables 1-3 for "COSMOS: COmmunity and Single Microbe Optimisation System": Supplementary Figures.docx

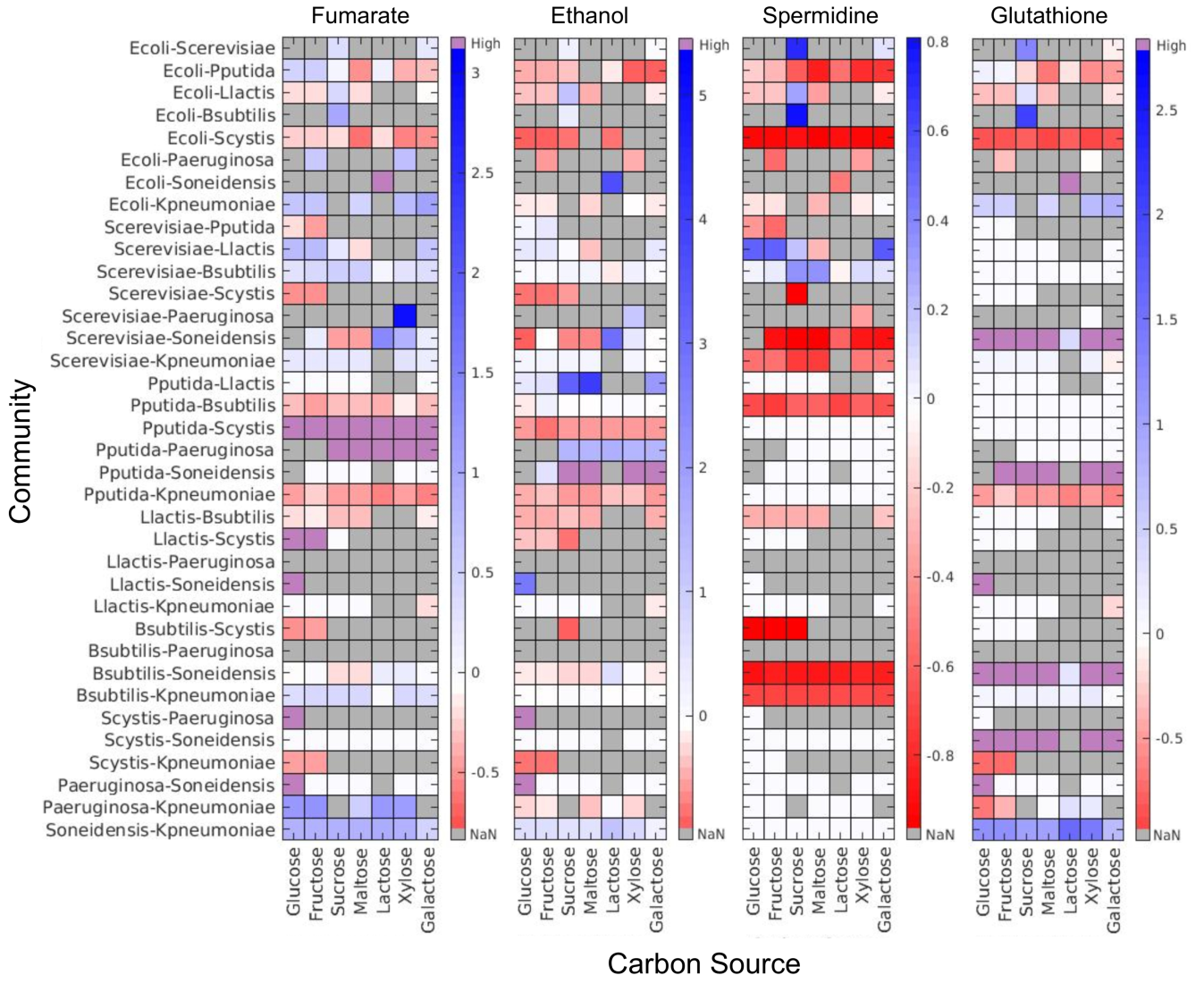

**Suppl. Fig 1. Effect of carbon source on the biosynthetic capability of different products in communities vs monocultures.** This figure illustrates the productivity ratio of specific products across different communities under varying carbon sources in an aerobic-rich environment. Positive values (blue) indicate an increase in the productivity of a given product in the community compared to the monocultures, while negative values (red) reflect a decline in productivity. If the product is only produced by the community and not by the monocultures, the value is shown in purple and labelled as 'High.' Conversely, when the community fails to grow on a specific carbon source, the value is represented in grey and marked as 'NaN.'
