## Supplementary Tables 1-3 for "COSMOS: COmmunity and Single Microbe Optimisation System": Supplementary Tables.docx

**Contents**

Supplementary Table 1 – List of organisms

Supplementary Table 2 – Minimal Medium composition

Supplementary Table 3 – Rich Medium composition

Supplementary Table 4 – List of products

Supplementary Table 5 – Best microbial system under different environments across four products

Supplementary Table 6 – Best microbial system under different environments across all products

**Supplementary Table 1. List of organisms**

| **Organism** | **Aerobic/Anaerobic** |
| --- | --- |
| *Escherichia coli* | Both |
| *Saccharomyces cerevisiae* | Both |
| *Pseudomonas putida* | Obligate aerobe |
| *Lactococcus lactis* | Both |
| *Bacillus subtilis* | Both |
| *Synechocystis spp.* | Both |
| *Pseudomonas aeruginosa* | Both |
| *Shewenella oneidensis* | Both |
| *Klebsiella pneumoniae* | Both |
| *Clostridium ljungdahlii* | Obligate anaerobe |

**Supplementary Table 2. Minimal Medium composition**

| **Minimal medium** | |
| --- | --- |
| Glucose | Oxygen |
| 4 Aminobenzoate | Phosphate |
| Biotin | Pantothenate |
| Calcium | Riboflavin |
| Cobalamin | Sulphate |
| Chlorine | Zinc |
| Cobalt | Selenium |
| Copper(II) | Thiamine |
| Iron(II) | Pyridoxal |
| Iron(III) | Hydrogen sulphide |
| Folate | Uracil |
| Hydrogen | L-alanine |
| Water | L-glutamate |
| Potassium | L-leucine |
| Magnesium | L-threonine |
| Manganese | L-valine |
| Molybdenum | L-isoleucine |
| Sodium | L-arginine |
| Nicotinamide | L-serine |
| Ammonium | Nicotinate |
| Nickel | Sulphite |
| Nitrate |  |

**Supplementary Table 3. Rich Medium composition**

| **Rich Medium** | |
| --- | --- |
| Glucose | L-alanine |
| 4 Aminobenzoate | L-glutamate |
| Biotin | L-leucine |
| Calcium | L-threonine |
| Cobalamin | L-valine |
| Chlorine | L-isoleucine |
| Cobalt | L-arginine |
| Copper(II) | L-serine |
| Iron(II) | Nicotinate |
| Iron(III) | Sulphite |
| Folate | L-aspartate |
| Hydrogen | L-asparagine |
| Water | L-cysteine |
| Potassium | L-glutamine |
| Magnesium | L-glycine |
| Manganese | L-histidine |
| Molybdenum | L-lysine |
| Sodium | L-methionine |
| Nicotinamide | L-phenylalanine |
| Ammonium | L-proline |
| Nickel | L-tryptophan |
| Nitrate | L-tyrosine |
| Oxygen | Adenosylcobalamin |
| Phosphate | Guanine |
| Pantothenate | Orotate |
| Riboflavin | Xanthine |
| Sulphate | Lead |
| Zinc | Protoheme |
| Selenium | Pimelate |
| Thiamine | Choline |
| Pyridoxal | Thymidine |
| Hydrogen sulphide | Carbon Dioxide |
| Uracil |  |

**Supplementary Table 4. List of products**

| **#** | **Product** | **Classification** |
| --- | --- | --- |
| 1 | Succinate | Organic acid |
| 2 | Pyruvate |  |
| 3 | Formate |  |
| 4 | L-Lactate |  |
| 5 | D-Lactate |  |
| 6 | Acetate |  |
| 7 | Fumarate |  |
| 8 | Gluconate |  |
| 9 | Propionate | Carboxylic acid |
| 10 | Adipic acid |  |
| 11 | Sorbitol | Sugar alcohol |
| 12 | Xylitol |  |
| 13 | Ethanol | Alcohol |
| 14 | Methanol |  |
| 15 | Butanol |  |
| 16 | Propane-1,2-diol | Diol |
| 17 | Propane-1,3-diol |  |
| 18 | 2,3 Butanediol |  |
| 19 | Glycerol | Triol |
| 20 | Hydrogen | Gas |
| 21 | Butyrate | Scfa |
| 22 | Spermidine | Polyamine |
| 23 | Putrescine |  |
| 24 | Catechol | Phenol |
| 25 | Glutathione | Tripeptide |

**Supplementary Table 5. Best microbial system under different environments across four products**

| **Aerobic Rich Environment** | | | | | | | | |
| --- | --- | --- | --- | --- | --- | --- | --- | --- |
| **#** | **Fumarate** | | **Ethanol** | | **Spermidine** | | **Glutathione** | |
|  | **Microbial system** | **Productivity (mmol/L/hr)** | **Microbial system** | **Productivity (mmol/L/hr)** | **Microbial system** | **Productivity (mmol/L/hr)** | **Microbial system** | **Productivity (mmol/L/hr)** |
| 1 | *P. aeruginosa - K. pneumoniae* | 1.27 | *S. oneidensis - K. pneumoniae* | 2.04 | *E. coli* | 0.07 | *S. oneidensis - K. pneumoniae* | 0.26 |
| 2 | *E. coli - K. pneumoniae* | 1.21 | *S. cerevisiae - K. pneumoniae* | 1.42 | *B. subtilis* | 0.05 | *E. coli - K. pneumoniae* | 0.25 |
| 3 | *S. oneidensis - K. pneumoniae* | 1.10 | *K. pneumoniae* | 1.29 | *S. cerevisiae - L. lactis* | 0.02 | *E. coli* - *P. putida* | 0.19 |
| 4 | *E. coli* - *P. putida* | 1.07 | *E. coli* | 1.10 | *S. cerevisiae* | 0.03 | *E. coli* | 0.17 |
| 5 | *B. subtilis - K. pneumoniae* | 0.88 | *P. aeruginosa* - *S. oneidensis* | 1.04 |  | 0.02 | *B. subtilis - K. pneumoniae* | 0.13 |
| **Aerobic Minimal Environment** | | | | | | | | |
| **#** | **Fumarate** | | **Ethanol** | | **Spermidine** | | **Glutathione** | |
|  | **Microbial system** | **Productivity (mmol/L/hr)** | **Microbial system** | **Productivity (mmol/L/hr)** | **Microbial system** | **Productivity (mmol/L/hr)** | **Microbial system** | **Productivity (mmol/L/hr)** |
| 1 | *P. aeruginosa - K. pneumoniae* | 0.58 | *S. oneidensis - K. pneumoniae* | 1.12 | *S. cerevisiae - B. subtilis* | 0.029 | *E. coli - K. pneumoniae* | 0.06 |
| 2 | *E. coli - K. pneumoniae* | 0.50 | *S. cerevisiae - P. aeruginosa* | 0.73 | *E. coli - P. aeruginosa* | 0.026 | *E. coli - P. putida* | 0.043 |
| 3 | *S. cerevisiae - P. aeruginosa* | 0.47 | *P. putida* - *S. oneidensis* | 0.73 | *E. coli - S. cerevisiae* | 0.024 | *S. oneidensis - K. pneumoniae* | 0.042 |
| 4 | *E. coli - P. aeruginosa* | 0.46 | *P. aeruginosa - K. pneumoniae* | 0.69 | *E. coli - P. putida* | 0.022 | *K. pneumoniae* | 0.038 |
| 5 | *Synechocystis - P. aeruginosa* | 0.38 | *E. coli - S. cerevisiae* | 0.65 | *E. coli - L. lactis* | 0.019 | *E. coli - P. aeruginosa* | 0.030 |
| **Anaerobic Rich Environment** | | | | | | | | |
| **#** | **Fumarate** | | **Ethanol** | | **Spermidine** | | **Glutathione** | |
|  | **Microbial system** | **Productivity (mmol/L/hr)** | **Microbial system** | **Productivity (mmol/L/hr)** | **Microbial system** | **Productivity (mmol/L/hr)** | **Microbial system** | **Productivity (mmol/L/hr)** |
| 1 | *E. coli* - *S. oneidensis* | 1.03 | *E. coli - S. oneidensis* | 1.37 | *E. coli - S. oneidensis* | 0.07 | *E. coli - S. oneidensis* | 0.26 |
| 2 | *E. coli - K. pneumoniae* | 0.89 | *C.* ljungdahlii - K*. pneumoniae* | 0.97 | *E. coli - K. pneumoniae* | 0.06 | *E. coli - K. pneumoniae* | 0.16 |
| 3 | *P. aeruginosa* - *S. oneidensis* | 0.33 | *S. oneidensis - K. pneumoniae* | 0.79 | *C. ljungdahlii* - *B. subtilis* | 0.02 | *S. oneidensis - K. pneumoniae* | 0.10 |
| 4 | *S. oneidensis - K. pneumoniae* | 0.33 | *C. ljungdahlii* - *B. subtilis* | 0.76 | *E. coli* | 0.02 | *P. aeruginosa* - *S. oneidensis* | 0.07 |
| 5 | *B. subtilis - K. pneumoniae* | 0.29 | *P. aeruginosa* - *S. oneidensis* | 0.68 | *B. subtilis* | 0.01 | *E. coli* | 0.05 |
| **Anaerobic Minimal Environment** | | | | | | | | |
| **#** | **Fumarate** | | **Ethanol** | | **Spermidine** | | **Glutathione** | |
|  | **Microbial system** | **Productivity (mmol/L/hr)** | **Microbial system** | **Productivity (mmol/L/hr)** | **Microbial system** | **Productivity (mmol/L/hr)** | **Microbial system** | **Productivity (mmol/L/hr)** |
| 1 | *P. aeruginosa - K. pneumoniae* | 0.34 | *P. aeruginosa* - *S. oneidensis* | 0.54 | *E. coli - P. aeruginosa* | 0.024 | *E. coli - P. aeruginosa* | 0.04 |
| 2 | *E. coli - P. aeruginosa* | 0.31 | *P. aeruginosa - K. pneumoniae* | 0.45 | *E. coli - C. ljungdahlii* | 0.0049 | *P. aeruginosa - K. pneumoniae* | 0.02 |
| 3 | *P. aeruginosa* - *S. oneidensis* | 0.20 | *E. coli - C. ljungdahlii* | 0.38 | *C. ljungdahlii* - *B. subtilis* | 0.0047 | *S. oneidensis - K. pneumoniae* | 0.02 |
| 4 | *B. subtilis - K. pneumoniae* | 0.10 | *E. coli - P. aeruginosa* | 0.36 | *B. subtilis* | 0.0041 | *P. aeruginosa* - *S. oneidensis* | 0.01 |
| 5 | *S. oneidensis - K. pneumoniae* | 0.09 | *E. coli - S. oneidensis* | 0.34 | *E. coli* | 0.0027 | *B. subtilis - K. pneumoniae* | 0.01 |

**Supplementary Table 6. Best microbial system under different environments across all products**

| **#** | **Aerobic-rich** | | **Aerobic-minimal** | | **Anaerobic-rich** | | **Anaerobic-minimal** | |
| --- | --- | --- | --- | --- | --- | --- | --- | --- |
|  | **Microbial system** | **No. of Products** | **Microbial system** | **No. of Products** | **Microbial system** | **No. of Products** | **Microbial system** | **No. of Products** |
| 1 | *E. coli* | 16 | *E. coli* | 15 | *E. coli* | 16 | *E. coli - P. aeruginosa* | 16 |
| 2 | *K. pneumoniae* | 14 | *K. pneumoniae* | 14 | *E. coli - K. pneumoniae* | 16 | *E. coli* | 15 |
| 3 | *S. cerevisiae* | 14 | *S. cerevisiae - L. lactis* | 14 | *E. coli - S. oneidensis* | 16 | *B. subtilis - K. pneumoniae* | 14 |
| 4 | *S. cerevisiae - L. lactis* | 14 | *E. coli - P. aeruginosa* | 13 | *K. pneumoniae* | 13 | *E. coli - C. ljungdahlii* | 14 |
| 5 | *S. cerevisiae - K. pneumoniae* | 13 | *E. coli - K. pneumoniae* | 12 | *S. oneidensis - K. pneumoniae* | 13 | *P. aeruginosa - K. pneumoniae* | 14 |
| 6 | *S. oneidensis - K. pneumoniae* | 13 | *S. cerevisiae* | 12 | *B. subtilis* | 11 | *S. oneidensis - K. pneumoniae* | 13 |
| 7 | *B. subtilis* | 12 | *S. cerevisiae - P. aeruginosa* | 12 | *P. aeruginosa - S. oneidensis* | 11 | *K. pneumoniae* | 12 |
| 8 | *B. subtilis - K. pneumoniae* | 11 | *S. oneidensis - K. pneumoniae* | 12 | *B. subtilis - K. pneumoniae* | 9 | *P. aeruginosa - S. oneidensis* | 11 |
| 9 | *E. coli - K. pneumoniae* | 11 | *B. subtilis* | 11 | *C. ljungdahlii - B. subtilis* | 8 | *C. ljungdahlii - B. subtilis* | 10 |
| 10 | *P. aeruginosa - S. oneidensis* | 11 | *E. coli - P. putida* | 10 | *L. lactis - S. oneidensis* | 8 | *E. coli - S. oneidensis* | 10 |
